# MVSE: an R-package that estimates a climate-driven mosquito-borne viral suitability index

**DOI:** 10.1101/360701

**Authors:** Uri Obolski, Pablo N Perez, Christian J Villabona-Arenas, Nuno R Faria, José Lourenço

## Abstract

**Background:** Viruses such as dengue, Zika, yellow fever and chikungunya depend on mosquitoes for transmission. Their epidemics typically present periodic patterns, linked to the underlying mosquito population dynamics, which are known to be driven by natural climate fluctuations. Understanding how climate dictates the timing and potential of viral transmission is essential for preparedness of public health systems and design of control strategies.

**Results:** We developed the **M**osquito-borne **V**iral **S**uitability **E**stimator (MVSE) software package for the R programming environment. The package estimates a suitability index based on a climate-driven mathematical expression for the basic reproductive number (R0) of a well established mathematical model for the transmission dynamics of mosquito-borne viruses. By accounting for local humidity and temperature, as well as viral, vector and human priors, suitability can be estimated for specific host and viral species, as well as different regions of the globe.

**Conclusions:** Here, we describe the background theory and biological interpretation of the new suitability index, as well as the implementation, basic functionality, research and educational potentials of the MVSE R-package. The package is freely available under the GPL 3.0 license.

## Introduction

Common mosquito-borne viruses include the dengue (DENV), chikungunya (CHIKV), Zika (ZIKV), yellow fever (YFV), Rift Valley fever (RVFV), West-Nile (WNV) and Japanese encephalitis (JEV) viruses. Due to ongoing human and climatic trends that favour the establishment of mosquitoes and movement of infectious hosts, these pathogens are becoming increasingly detrimental for human public health (Jaenisch et al., 2014; Schwarz et al., 2012; Stanaway et al., 2016). For instance, recent outbreaks of ZIKV and CHIKV (2014–2017) have taken a toll on populations of multiple island-nations (Cauchemez et al., 2016; Duffy et al., 2009; Lourenço et al., 2018), South America and the Caribbean (Faria et al., 2016, 2017b; Lourenço et al., 2017a; Rodrigues Faria et al., 2016). In Africa, YFV outbreaks in Angola and other countries have recently (2016–2017) developed into a global vaccination crisis (Kraemer et al., 2016; Wu et al., 2016). More recently (2017–2018), YFV has also emerged from its sylvatic cycle in Brazil (Faria et al., 2018). Not only is the absolute scale of these outbreaks unprecedented, but the emergence of severe pathologies such as ZIKV-related neonatal microcephaly, CHIKV-induced Guillain-Barré syndrome and a very high YFV mortality rate have had substantial negative socio-economic impacts (Cao-Lormeau et al., 2016; Johansson et al., 2016; Oehler et al., 2015; de Oliveira et al., 2017).

The evolutionary and host-pathogen history of mosquito-borne viruses is vastly diverse. For instance, while a strong ecological barrier separates human and zoonotic variants of DENV and ZIKV (Holmes and Twiddy, 2003; Lessler et al., 2016), such a barrier is less evident for YFV, WNV and JEV (Lanciotti, 1999; Rosen, 1986). DENV also uniquely presents four antigenically distinct lineages (serotypes DENV1-4) with complex immunological responses (Flasche et al., 2016; Lourenço and Recker, 2016; Lourenço et al., 2017b), whereas WNV has the vastest host tropism including primates, equines, birds and reptiles (Mackenzie et al., 2004; Rosen, 1986). Importantly, however, the population biology of these viruses shares one unifying characteristic: the dynamics of their epidemic behaviour are inherently linked to the underlying population dynamics of their vector species. Mosquito-population dynamics are known to be dictated by a wide range of factors, such as climate, altitude, population density of humans or other mammals, pollution levels, and natural or artificial water reservoirs (Chen et al., 2012; Lozano-Fuentes et al., 2012; Monteiro et al., 2007; Williams et al., 2010). While most of these factors can dictate absolute population sizes (carrying capacity), seasonal oscillations in size are directly and indirectly driven by natural climate variations (Brady et al., 2013; Dutta et al., 2010; Honório et al., 2009; LaBeaud et al., 2011; Madeira et al., 2002; Mohammed and Chadee, 2011; Oo et al., 2011; Yasuno and Tonn, 1970).

Recently, we have developed a climate-driven, mathematical model of mosquito-borne viral transmission based on ordinary differential equations, which has been used to study the entomological and epidemiological determinants of the 2012 dengue outbreak in the island of Madeira (Portugal) (Lourenço and Recker, 2014), the 2014 dengue outbreak in Rio de Janeiro (Brazil) (Faria et al., 2017a), and the 2015–2017 ZIKV outbreak in Feira de Santana (Brazil) (Lourenço et al., 2017a). In these case studies, the model was fit to notified epidemic curves using Markov chain Monte Carlo, allowing the estimation of epidemiological parameters such as the basic reproduction number (R0) and the effective reproduction number (Re). The success of this framework stems from its data-driven approach, including temporal climatic series used to parameterise both vector and viral variables under mathematical relationships derived from experimental studies. The framework includes estimation of the effect of local climatic variables on viral and vector parameters, which enables it to be generalized to regions with different climates. Furthermore, by allowing specific vector, viral and host parametrization, this framework is also general enough to be applied to different host-pathogen systems.

In this article, we translate our experience with this modelling framework into the MVSE R-package, which is capable of estimating and analysing a *mosquito-borne viral suitability index*. Given local climatic variables, this index can be used as a proxy for the timing and scale of transmission potential, and is applicable to different viruses, vectors and regions. The main goal of this package is to offer a software tool free of the mathematical complexity of epidemiological models and of the need to depend on epidemiological time series (incidence) of the virus under study. Here, we describe in detail the package’s functionalities using real world examples and argue for its public health and academic potential.

## Theory, Design and Implementation

Recently, we have shown that a suitability measure called *index P* can be derived from the formula of R0 without the need to fit epidemic curves, as long as the relationships of some of R0’s parameters with climate are known. The index P was shown to be correlated with the timing and scale of both local mosquito infestation measures and dengue notifications in Myanmar (Pn et al., 2017). Here, we describe the rationale behind the index P in detail, and demonstrate its potential in other epidemiological contexts.

### The suitability index P

#### Derivation and interpretation

The transmission potential of a pathogen can be summarized through the basic (R0) and effective (Re) reproduction numbers. The R0 of a pathogen is the number of secondary cases generated, on average, by a single infected host in a totally susceptible population. In the case of mosquito-borne viruses, R0 can also be interpreted as the sum of the reproductive potential (transmission) of each adult female mosquito, *P*_(_*_u,t_*_)_, over the total number of female mosquitoes per human, M, in a totally susceptible host population (equation 1). The Re of a mosquito-borne virus can be interpreted in a similar manner, but taking into consideration the presence of immune hosts hampering transmission potential (equation 2, with *S_h_*, *S_v_* the proportion of susceptible humans and mosquitoes, respectively).

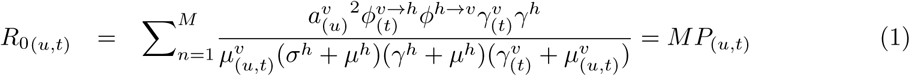

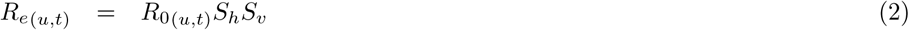

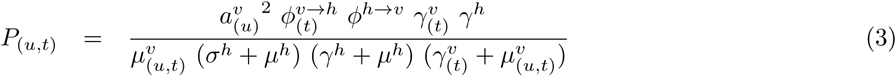

There are a total of eight parameters in the expression of R0, four of which are climate-independent (the human life-span 1/*μ^h^*, the transmission probability from infected human to mosquito per bite ϕ^*p→v*^, the human infectious period 1/*σ^h^*, and human incubation period 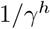, and four are climate-dependent (the life-span of adult mosquitoes 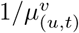, the extrinsic incubation period 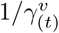, the daily biting rate 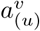 and the probability of transmission from infected mosquito to human per bite 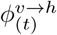. Climate-dependent parameters are defined as functions dependent on humidity (u) and temperature (t), which have previously been determined in experimental studies through laboratory estimates of entomological data under various climate conditions. A list of these functions, sources and details can be found in the Supporting Information **S2 Text**.

Quantification of R0 (and Re) requires an estimation of the number of adult female mosquitoes per human (M). It is rarely the case that adequate estimations of M exist for regions or mosquito-species of interest. Given that the mosquito population size fluctuates in and out of season, M is expected to present oscillatory behaviour. Thus, the R0 (and Re) of a mosquito-borne virus, dependent on M, also presents seasonal oscillations. However, such oscillations are only partially driven by M, and are also determined by *P*_(_*_u,t_*_)_ (equation 1). In our mathematical framework, *P*_(_*_u,t_*_)_'s contribution to this oscillatory behaviour stems from the four climate-driven parameters: the life-span of adult mosquitoes, the extrinsic incubation period, the daily biting rate, and the probability of transmission from infected mosquito to human per bite.

Theoretically, the potential for outbreaks is determined by the epidemic thresholds of R0>1 or Re>1. *P*_(_*_u,t_*_)_ is a positive number that can be interpreted as the absolute potential of an adult female mosquito. Thus, if at least one female mosquito exists per human (*M* >= 1) and *P*_(_*_u,t_*_)_ *>* 1, then *R*0 = *MP*_(_*_u,t_*_)_ > 1 and epidemic growth is possible. However, interpreting *P*_(_*_u,t_*_)_ on its own means that no direct assessment on the classic epidemic thresholds can be made. We thus do not make particular interpretations of *P*_(_*_u,t_*_)_ > 1, but instead argue and show that its absolute value is informative for transmission potential when assessed locally in time or between regions.

*P*_(_*_u,t_*_)_ can be estimated for any region for which humidity and temperature are available, and can be parameterized for any species of virus, host or vector. We thus define *P*_(_*_u,t_*_)_ as the *mosquito-borne viral suitability index P*, using a complementary terminology to existing *vector suitability* indices (Gardner et al., 2018; Kraemer et al., 2015a,b). A key difference from *vector suitability* indices is that the numerical scale of the index P has direct biological interpretation: P is the reproductive (transmission) potential of an adult female mosquito.

#### Estimation

In previous studies on ZIKV and DENV, we estimated R0 and Re by fitting a dynamic transmission model to notified epidemic curves within a Bayesian Markov chain Monte Carlo (bMCMC) framework, whereby unobserved parameters were estimated by their resulting posterior distributions (Faria et al., 2017a; Lourenço et al., 2017a; Lourenço and Recker, 2014). Here, however, our goal is to estimate suitability for transmission, through *P*_(_*_u,t_*_)_, independently of the availability of epidemiological data.

The Bayesian strategy implemented in MVSE focuses on defining priors for all the parameters in the expression of *P*_(_*_u,t_*_)_ for which adequate support exists in the scientific literature. For the climate-driven parameters (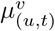, 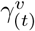, 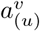 and 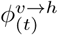), which affect the amplitude and timing of transmission, such informed priors bound the possible solutions of *P*_(_*_u,t_*_)_ for the virus, mosquito and region of interest. Methodological details, MVSE code and examples of the estimation steps of *P*_(_*_u,t_*_)_ can be found in Supporting Information **S2 Text**.

## Selection of Results

### Index P dynamics in Recife and São Paulo

We first present a comparison of the estimated index P per day in two major urban centres - Recife and São Paulo (Brazil) - for the time period between 2005 and 2016. Recife is located in the state of Pernambuco in the Northeast of Brazil with a tropical savanna climate, while São Paulo is located in the south with a humid subtropical climate (Köppen climate classification (Peel et al., 2007)). The two cities are known to have contrasting epidemic and endemic histories for mosquito-borne viruses. For instance, Recife has experienced a large ZIKV epidemic and is known to be endemic to DENV and CHIKV (de Araújo et al., 2016; Magalhaes et al., 2017; Sabino et al., 2016). In contrast, São Paulo is yet to experience a sustained ZIKV epidemic, DENV incidence is known to be low for the national average (Teixeira et al., 2013), and transmission of mosquito-borne viruses is unstable, with phylogenetic studies showing that epidemics are seeded by new lineages almost every year (Faria et al., 2017a,b). In the notified dengue data that we obtained for 2007–2012 (**S3 Table**), the total incidence per 100k individuals in Recife was approximately 11-fold higher than it was in São Paulo (2184.3 notifications versus 195.2).

The input (climate time series, user-defined priors) and index P output for the two cities is summarized in Figure 1 (for a more complete set of possible outputs see **S1 Text**). Here, for simplicity, and for a lack of evidence of significant differences between the two cities, we assume all informed priors to be the same (Figures 1 b,d). As expected, differences can be seen in the local climatic trends, with São Paulo showing, for instance, lower temperatures compared to Recife (Figures 1 a,c). Figures 1 e,f show the estimated index P for the two cities, which present significantly different dynamics. For instance, the mean index P in Recife is estimated to be 1.5-fold higher than in São Paulo (1.26 versus 0.84). Seasonality is also different, with Recife showing less pronounced oscillations and with mean P generally maintained above 1 (i.e. one female mosquito per human would necessarily be enough for persistent transmission, equation 1). In contrast, São Paulo shows pronounced oscillations, with P presenting substantial troughs.

**Fig 1.**
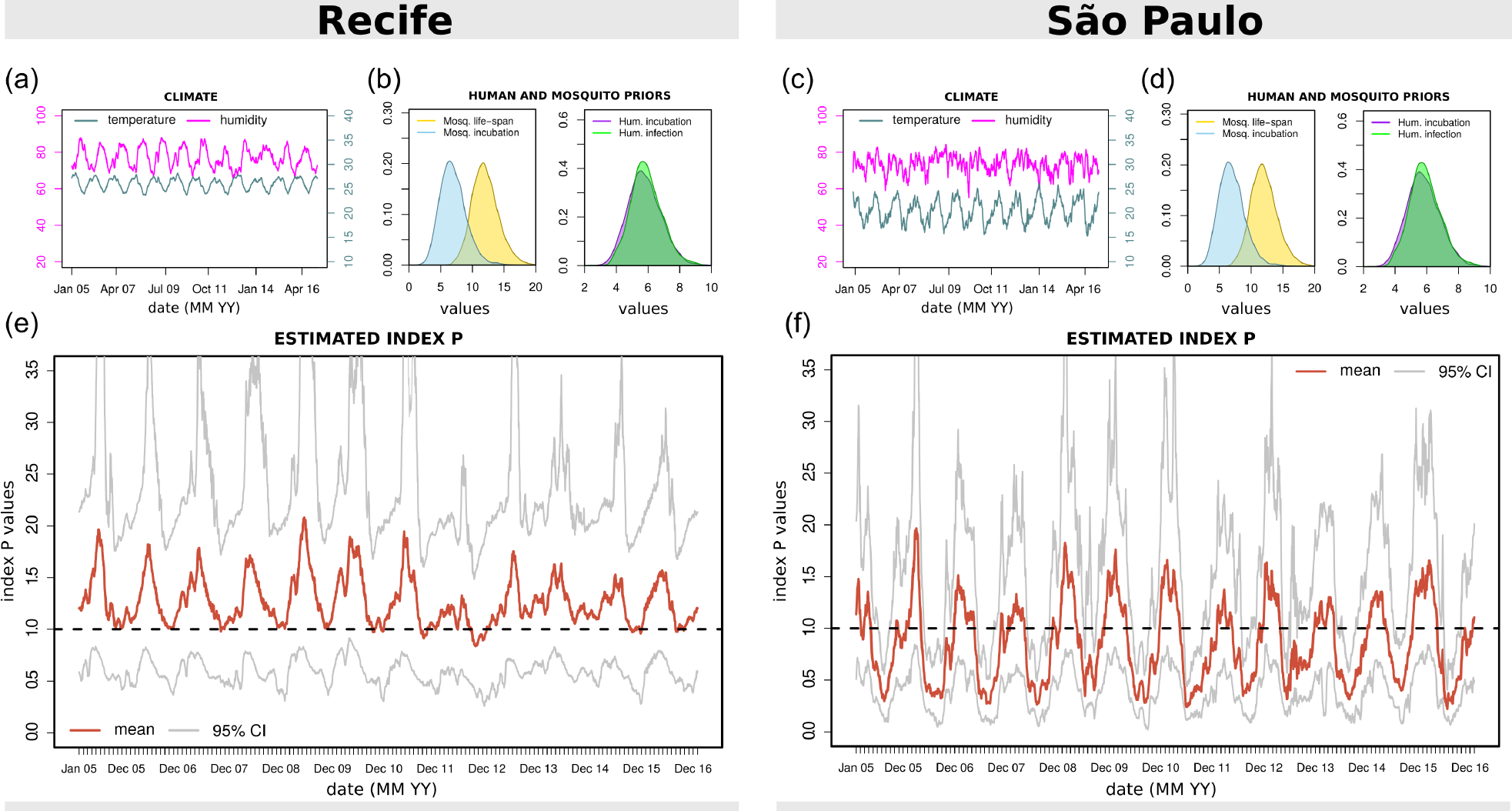
Climate, priors and estimated index P for Recife and São Paulo. (**a,c**) Local humidity (magenta) and temperature (turquoise) time series per day. (**b,d**) Examples of MVSE informed priors: mosquito life-span (blue) and incubation period (yellow); human incubation (purple) and infectious (green) periods. (**e,f**) Estimated index P per day with mean (red) and 95% confidence interval (grey). Priors were assumed to be the same for Recife and São Paulo. See Supplementary Information **S2 Text** for prior distributions.

These key differences in index P are in accordance to what is known about the transmission potential of mosquito-borne viruses in the two cities. For instance, the stabler seasonality patterns and higher index P in Recife suggests year-round potential for single adult mosquitoes to contribute to epidemic expansion; and can thus help explain why dengue incidence in Recife can be one order of magnitude higher than in São Paulo. Concurrently, the substantial troughs of index P estimated in São Paulo suggest very low transmission potential off-season, which may partially explain why persistence of mosquito-borne viral lineages between seasons is rarely observed in the city.

### Index P seasonality in Recife and São Paulo

We next present two visual outputs from MVSE, comparing some of the seasonal differences in the estimated index P between Recife and São Paulo (for a more complete set of possible outputs see **S1 Text**). Figure 2 shows the timing of highest index P and sensitivity of P to each of the climatic variables. For visual convenience, we restrict MVSE’s output to the period of four years, between 2010 and 2013.

**Fig 2.**
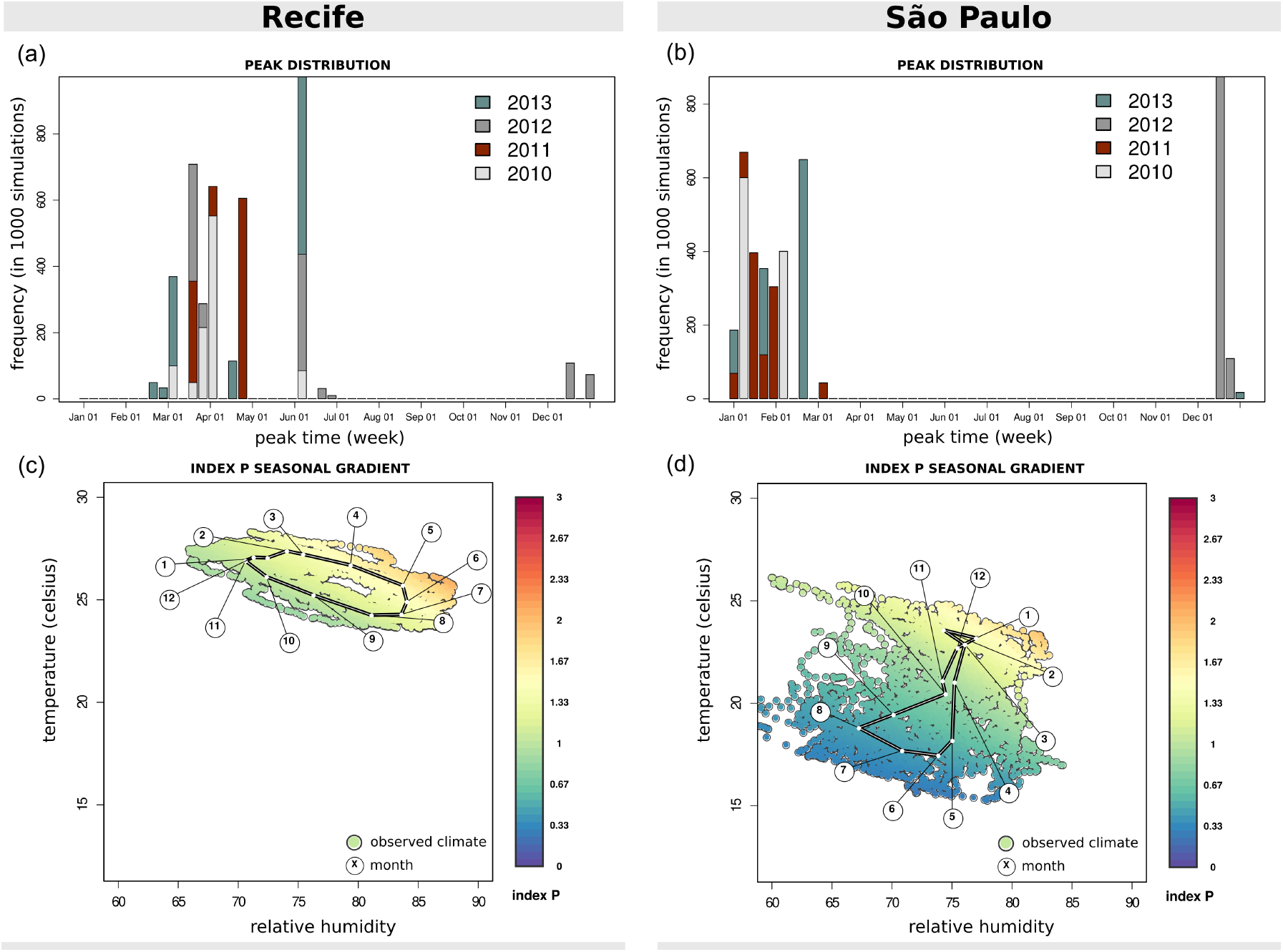
Seasonality of index P for Recife and São Paulo. (**a,b**) For each year colored differently, the week with highest suitability is identified across all estimated index P solutions of that year (solutions’ mean and 95% CI in Figure 1). The frequency of each peak week shown across the 1000 simulations. (**c,d**) Two dimensional sensitivity of mean index P per humidity (x) and temperature (y) observed time point (day). Each point is a combination of observed climate variables, colored according to the mean index P estimated (color scale on the right). The white dots (over the black link) mark the mean humidity and temperature of each month over the period of the data (2005–2016); the floating circles with numbers identify each month’s number.

For each year and city, the week with highest suitability is identified across all estimated index P solutions *(simulations*, of which the mean and 95% CI is in Figures 1 e,f). As seen in Figures 2 a,b, the two cities vary in their detected peak weeks. In Recife, suitability tends to peak between March and May (regional autumn), with the occasional occurrence of higher suitability in June (in particular in the years 2012 and 2013). In contrast, suitability peaks earlier in São Paulo, between late December and early February (regional summer).

MVSE also offers visual output which highlights the regional sensitivity of the index P to climatic changes in time (Figures 2 c,d). Once again, the differences between the cities are clear. Humidity and temperature values are less variable in Recife than in São Paulo, and Recife has both higher humidity and temperature. This visualisation also reveals a clear gradual trend in suitability across the months in Recife. In contrast, this gradient is only clear for São Paulo in the winter months (May - September).

### Index P spatio-temporal characterization across Brazil

In the previous Figures we demonstrated how the index P can be used to compare transmission potential and timing between two Brazilian cities. Here, we demonstrate the usefulness of index P to characterize the transmission potential across vast geographical ranges, using Brazil as a case study. We use the WorldClim V2 data set, which holds mean worldwide climatic data per month, during 1970–2000 (see Data accessibility for details). Our analysis is performed over a regular grid of Brazil, in which each pixel represents an area of ≈ 340km^2^. For each pixel, we use 12 time points (months) of humidity and temperature to run MVSE and estimate the dynamics of index P over one year. Although computationally intensive, due to the size of the grid, this exercise only employs functions available in MVSE. For further details on the data, and outputs of a similar exercise over the entire South American continent, see Supplementary Information **S2 Text**. Figure 3 presents a series of maps derived from MVSE’s output.

**Fig 3.**
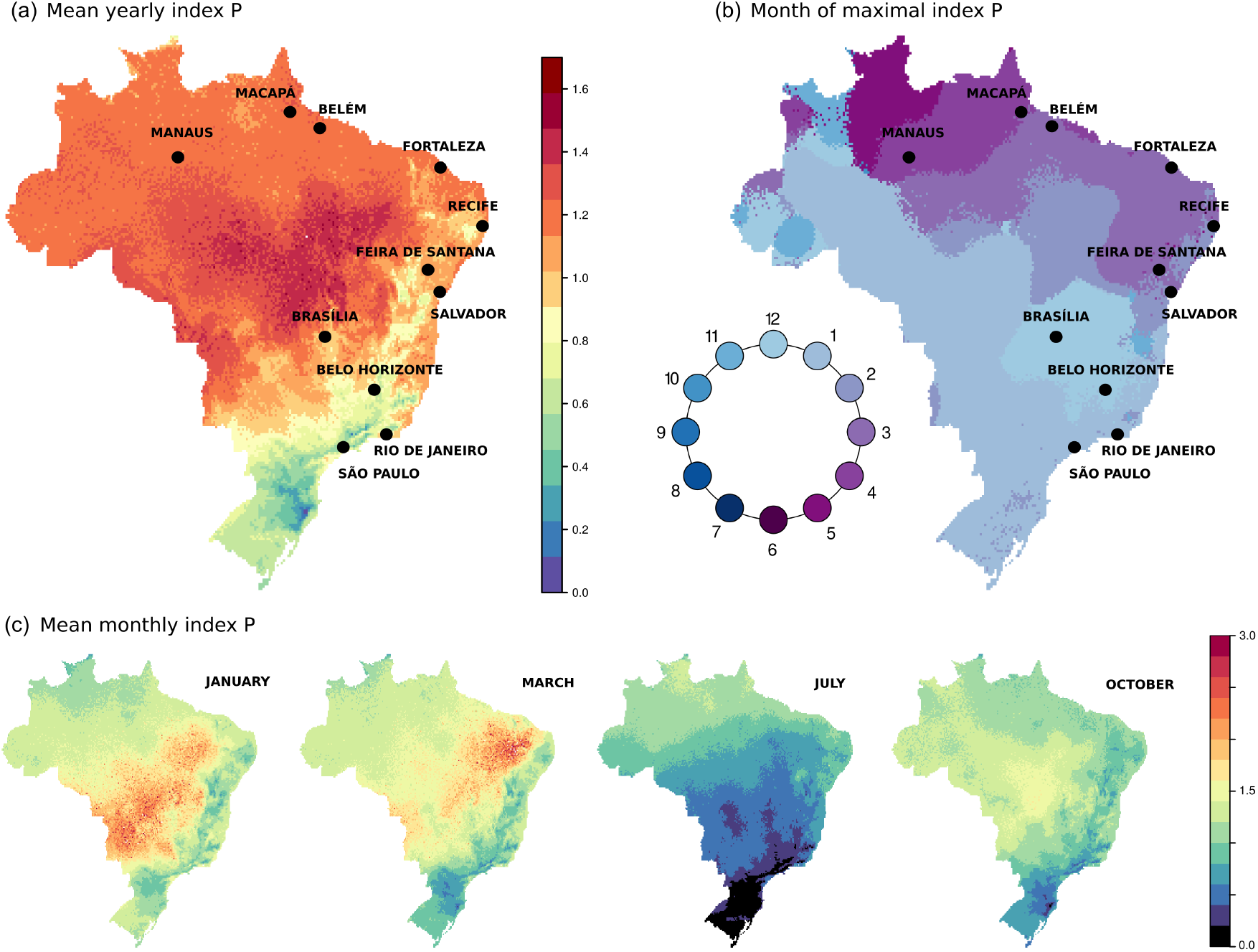
Spatio-temporal characterization of index P across Brazil. (**a**) Map presents the mean index P per pixel (≈ 340Km^2^). Values colored according to scale on the right. (**b**) Using the estimated index P of each pixel, with 12 points representing months, the month with highest index P is identified. Each pixel is colored according to that month, with the color scale represented in a circle. (**c**) Same as (a), but presenting the mean for selected months. In all maps, priors were assumed to be the same per pixel, as used for Recife and São Paulo. See Supplementary Information **S2 Text** for prior distributions. These solutions are made available as Supplementary Files.

The estimated mean suitability presents high variation across Brazil (Figure 3 a). As expected, the southern regions present the lowest transmission potential (colder colors), while the highest potential is estimated in the center and along the northern coast (warmer colors). The elevated regions of the country, running in parallel to the eastern coast, present a pattern of intermediate suitability.

We further identify the month of peak suitability for each region (Figure 3 b). This discretization highlights a seasonal gradient in the month of maximal suitability across Brazil. In the southern regions, peak suitability is during the early summer months, and moving north along the eastern and then northern coasts shifts peak suitability further into regional autumn. Notably, no region of Brazil is seen to experience peak suitability during the core winter months (July - September, darker blue).

We also map the mean monthly index P for a subset of months (Figure 3 c). In these maps, the generally lower suitability in the south is clearly highlighted, as is the fact that patches of higher suitability occur at different months in different regions.

### Index P and notified cases in Brazillian cities

In Figures 1 and 2 we characterised the index P for the cities of Recife and São Paulo. Here, we use climate data from the same time period for other cities in Brazil, and present the correlation between the estimated suitability and dengue notified cases. Notifications were per month, summed up for the period 2007–2012 (see Data accessibility for details). We estimated index P per day in the period 2005 to 2016, and then calculated mean P for each of the 12 months across the years. By averaging across 11 years, we effectively quantified suitability into what can be termed a *typical year*, i.e. the expected average monthly suitability of each city in any year.

Figure 4 presents the typical year suitability against the case notifications, per city. Pearson's correlation (*ρ*) shows that the index P is highly and positively correlated with dengue notifications in Brazilian cities. These results are similar to those we have previously demonstrated for Myanmar at the country and city level (Pn et al., 2017).

**Fig 4.**
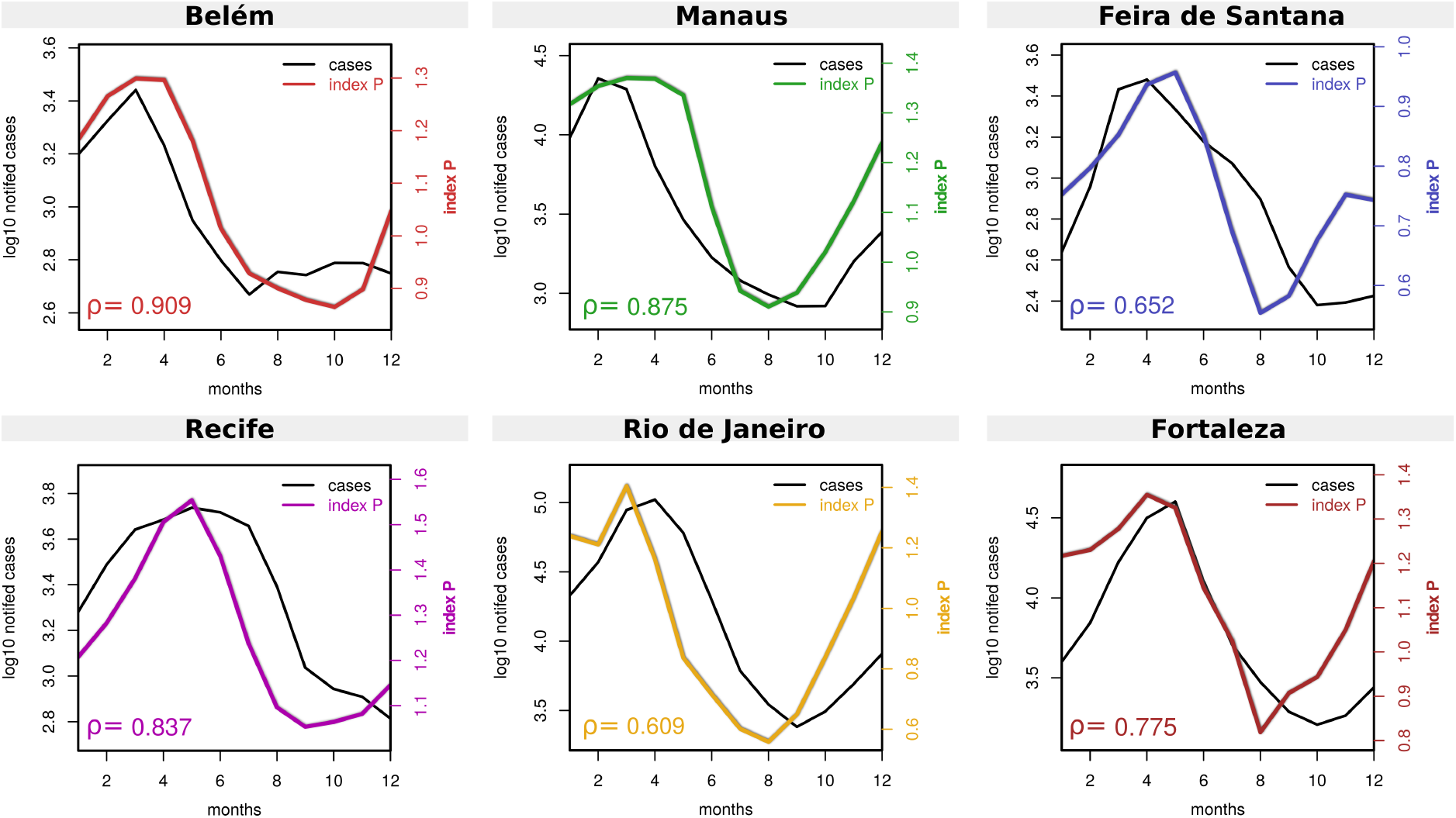
Correlation of index P and notified cases in Brazilian cities. Time series of index P (2005–2016) with dengue notified case data (2007–2012) is shown for 6 cities: Belém (red), Manaus (green), Feira de Santana (blue), Recife (purple), Rio de Janeiro (yellow) and Fortaleza (brown). Pearson’s correlation coefficient (*ρ*) is shown within each subplot. For a visual representation of linear regressions and p-values see Figure 5 in **S1 Text**.

### Conclusions and Future Directions

In this manuscript, we have introduced in detail a mosquito-borne viral suitability measure which we named index P. We apply and exemplify the capacities of index P to estimate transmission potential and characterize it in space and time using real world examples. We also introduce and provide MVSE, an R-package that implements various functionalities allowing to estimate and analyse the index P in regions for which humidity and temperature time series are available.

We believe that both MVSE and the index P are unique in the current literature. Other useful suitability indices exist, but these are based on complex approaches dependent on many variables from various sources, with no direct biological interpretation on the measured scale, and for which no freely available estimation tools exist. In contrast, index P is simply based on the R0 expression of a mosquito-borne dynamic model and two climate variables, has a direct biological interpretation, and is easily calculated through MVSE, a complete and freely available software tool.

With its aforementioned mathematical simplicity, dependency on only two climatic variables and freely available R-package MVSE, we foresee the potential of the index P not only for direct research purposes, but also for education (e.g. epidemiological courses on arboviruses) and field ento-epidemiology (e.g. entomologists and epidemiologists collecting real-time data). New ideas and methodological updates will be implemented in the MVSE package as the user base will keep growing. Future expansions of the package could include functionalities regarding the total number of female mosquitoes per human (M), or the inclusion of other local variables which may be related to the risk of mosquito-borne viruses (e.g. socio-economic variables, rainfall, mosquito control initiatives, etc.). Any future updates on the methodology and R-package will be made available in the package’s online repository (see **S1 Text**).

## Supporting information

**S1 Text.** Supplementary Figures. Examples of visual outputs produced by MVSE in the context of Recife and Saao Paulo that are not presented in the main text, and other figures that complement the main text.

**S2 Text.** Full methods and data description. Detailed description of package implementation, code examples with outputs, detailed methodology, estimation of index P across South America, spatial distribution of dengue notifications in Brazil, correlation of index P and dengue notifications in Brazil, and correlation of index P and dengue notifications in Myanmar.

**S1 Animation.** GIF file for South America. Animation with monthly index P estimations across South America.

**S2 Animation.** GIF file for Brazil. Animation with monthly index P estimations across Brazil.

**S1 Table.** Estimated index P per month across South America. Rdata file with the estimated index P across the continent. For details on how this data is organized, see Supplementary Information **S2 Text**.

**S2 Table.** Estimated index P per month across Brazil. Rdata file with the estimated index P across the country. For details on how this data is organized, see Supplementary Information **S2 Text**

**S3 Table.** Dengue notifications for Brazil. CSV file with total notified dengue cases per month summed over the period 2007–2012 per city (Manaus, Belem, Recife, Fortaleza, Feira de Santana, Rio de Janeiro).

**S4 Table.** MVSE solutions for entomological parameters in São Paulo. CSV file with entomological solutions (posteriors) for São Paulo.

**S5 Table.** MVSE solutions for index P in São Paulo. CSV file with mean, confidence intervals and smoothed index P for São Paulo.

**S6 Table.** Climate variables for São Paulo. CSV with date (time point), humidity and minimum temperature for São Paulo.

## Acknowledgments

UO received an EMBO postdoctoral fellowship. PNPG received travel and accommodation expenses from the Department of Global Health and Oriel College, University of Oxford. NRF is supported by a Royal Society and Wellcome Trust Sir Henry Dale Fellowship (204311/Z/16/Z), internal HEFCE GCRF grant 005073, and John Fell Research Fund Grant 005166. JL received funding from the European Research Council under the European Union’s Seventh Framework Programme (FP7/2007–2013, ERC grant number 268904 - DIVERSITY). CJVA was supported by an European Research Council Starting Grant (ERC-2017–STG, project ID: 757688). The funders had no role in study design, data collection and analysis, decision to publish, or preparation of the manuscript.

## Author’s contributions

JL developed the index P theory. PNP demonstrated the theory in the first case studies. JL, CJVA and NRF planned the development of the R-package. JL and UO curated the data, coded the R-package, performed the experiments and wrote the manuscript. All authors tested the code, and revised both the examples and manuscript text.

## Data accessibility

Climatic data (average relative humidity and minimum temperature) were collected from the Brazilian public repository ’Meteorological database for education and research’ (banco de dados meteorológicos para ensino e pesquisa) available at the BDMEP website for the cities of São Paulo, Recife, Manaus, Belém, Fortaleza, Feira de Santana and Rio de Janeiro (period 2005–2016). We make the São Paulo data available in **S6 Table**.

WorldClim V2 global data (Fick and Hijmans, 2017) were downloaded from a public repository online. Updated in July 2016, these data contain average monthly climatic measurements, representative of the period 1970–2000. The spatial resolution used was of 10 minutes (≈ 340*km*^2^). These data are not made available in this manuscript due to license restrictions, but are publicly available at WorldClim webpage. See Supporting Information **S2 Text** for more details.

Dengue notification data (by clinical evaluation, not molecularly confirmed) were obtained from the Brazilian ministry of health, available at the TABNET webportal. These notification data included the total number of dengue cases per month summed over the period 2007–2012 for the main municipality of each city (**S3 Table**).

